# Interdependent Iron and Phosphorus Availability Controls Photosynthesis Through Retrograde Signaling

**DOI:** 10.1101/2021.02.11.430802

**Authors:** Hye-In Nam, Zaigham Shahzad, Yanniv Dorone, Sophie Clowez, Kangmei Zhao, Nadia Bouain, Huikyong Cho, Seung Y. Rhee, Hatem Rouached

## Abstract

Iron deficiency hampers photosynthesis and is associated with chlorosis. We recently showed that iron deficiency-induced chlorosis depends on phosphorus availability. How plants integrate these cues to control chlorophyll accumulation is unknown. Here, we show that iron limitation downregulates photosynthesis genes in a phosphorus-dependent manner. Using transcriptomics and genome-wide association analysis, we identify two genes, a chloroplastic ascorbate transporter (*PHT4;4*) and a nuclear transcription factor (*bZIP58*), which prevent the downregulation of photosynthesis genes leading to the stay-green phenotype under iron-phosphorus deficiency. Joint limitation of these nutrients induces ascorbate accumulation by activating expression of an ascorbate biosynthesis gene, *VTC4*, which requires bZIP58. Exogenous ascorbate prevents iron deficiency-induced chlorosis in *vtc4* mutants, but not in *bzip58* or *pht4;4*. Our study demonstrates chloroplastic ascorbate transport is essential for preventing the downregulation of photosynthesis genes under iron-phosphorus combined deficiency. These findings uncover a molecular pathway coordinating chloroplast-nucleus communication to adapt photosynthesis to nutrient availability.

---

Chloroplasts are sites of photosynthesis, whose function requires numerous proteins encoded in the nuclear genome ^1^. Although plants tightly orchestrate chloroplast-to-nucleus signaling (retrograde control), it is poorly understood at the mechanistic level. Additionally, the adequate accumulation of nutrients such as iron (Fe) in chloroplasts is required for their optimal performance ^2,3^. Up to 80% of Fe in leaves is located in the chloroplasts^4,5^, where its ability to donate and accept electrons plays a central role in electron transfer reactions^6^. Fe is found in all electron transfer complexes PSI, PSII, cytochrome b6f complex, and ferredoxins and is required for the biogenesis of cofactors such as hemes and iron-sulfur clusters^7,8^. Plants grown under Fe-deficient (-Fe) environments show chlorotic symptoms^9^, and compromised photosynthesis ^2,3^. However, chlorotic leaves can also develop under high-phosphorus (P) conditions, despite replete Fe levels^10^, challenging the causal connection between Fe concentration and chlorophyll accumulation. Moreover, we recently reported that rice plants grown under a combined Fe and P deficiency (-Fe-P) do not exhibit a chlorotic phenotype^11^. These observations revealed a gap in our understanding of the interdependent effects of nutrient availability on regulating photosynthesis. Here, we addressed this issue through a combination of global gene expression analyses and genome-wide association studies (GWAS) to find expression quantitative trait loci (eQTLs) and uncovered a regulatory module that controls chlorophyll accumulation in response to Fe and P availability. This module involves an ascorbic acid (AsA) synthesis enzyme named VITAMINC4 (VTC4), a chloroplastic AsA transporter named PHOSPHATE TRANSPORTER 4;4 (PHT4;4), and a putative transcription factor named BASIC LEUCINE-ZIPPER 58 (bZIP58). The functioning of this module sheds light on the importance of chloroplast-nucleus communications under co-occurring nutrient deficiencies in controlling photosynthesis.

We previously reported that Fe deficiency-induced chlorosis depends on P availability in rice^11^. To investigate whether the interdependent effects of Fe and P availability on chlorosis are conserved across monocot and eudicot species, we phenotyped *Arabidopsis thaliana* Col-0 (eudicot) and *Lemna gibba* (monocot), along with *Oryza sativa* (monocot), under different regimes of Fe and P availability. Fe deficiency (-Fe) caused chlorosis in all three species, but only in the presence of P (-Fe+P) (Figure 1A-C). Quantification of chlorophyll content confirmed that -Fe significantly reduced the accumulation of chlorophyll in all three species (Figure 1D). However, under -Fe-P conditions, chlorophyll content was comparable to control (+Fe+P) conditions in these species (Figure 1D). Next, we focused on Arabidopsis to gain insights into the physiological and molecular processes underlying the recovery of chlorosis under -Fe-P conditions. First, we asked whether absence of chlorosis under -Fe-P is caused by an increase of Fe levels in shoots. Plants grown in -Fe+P conditions decreased total Fe in shoots by 2-fold compared to +Fe+P conditions (Figure 1E). On the other hand, under +Fe-P conditions, Fe levels increased by 2.2-fold relative to +Fe+P conditions (Figure 1E). Surprisingly, Fe levels in plants grown under -Fe-P were reduced and indistinguishable from the Fe levels in -Fe+P conditions (Figure 1E). Therefore, the lack of chlorosis under -Fe-P is unlikely to be caused by more Fe available in shoots. To further test this hypothesis, we assessed free Fe in leaves. Because FERRITIN1 (AtFER1) chelates Fe and its mRNA increases when Fe is in access, *AtFER1* gene expression can be used as a read-out for intracellular Fe nutritional status^12^. We thus quantified the expression of *AtFER1* in shoots under replete or deficient Fe and P in the growth media. Consistent with the total Fe levels, *AtFER1* expression was increased significantly under +Fe-P conditions relative to +Fe+P (Figure 1F). However, *AtFER1* expression decreased under both -Fe+P and -Fe-P conditions relative to +Fe+P (Figure 1F). Interestingly, -Fe-P condition caused a slightly bigger reduction in *AtFER1* expression than -Fe+P did, suggesting that there may be even less free Fe in -Fe-P conditions than in -Fe+P conditions. Taken together, these data show that the onset of chlorosis during -Fe requires sufficient P in the growth media, and that the “stay green” phenotype under the combined -Fe-P deficiency cannot be linked to Fe nutritional status in leaves.

**Figure 1.**
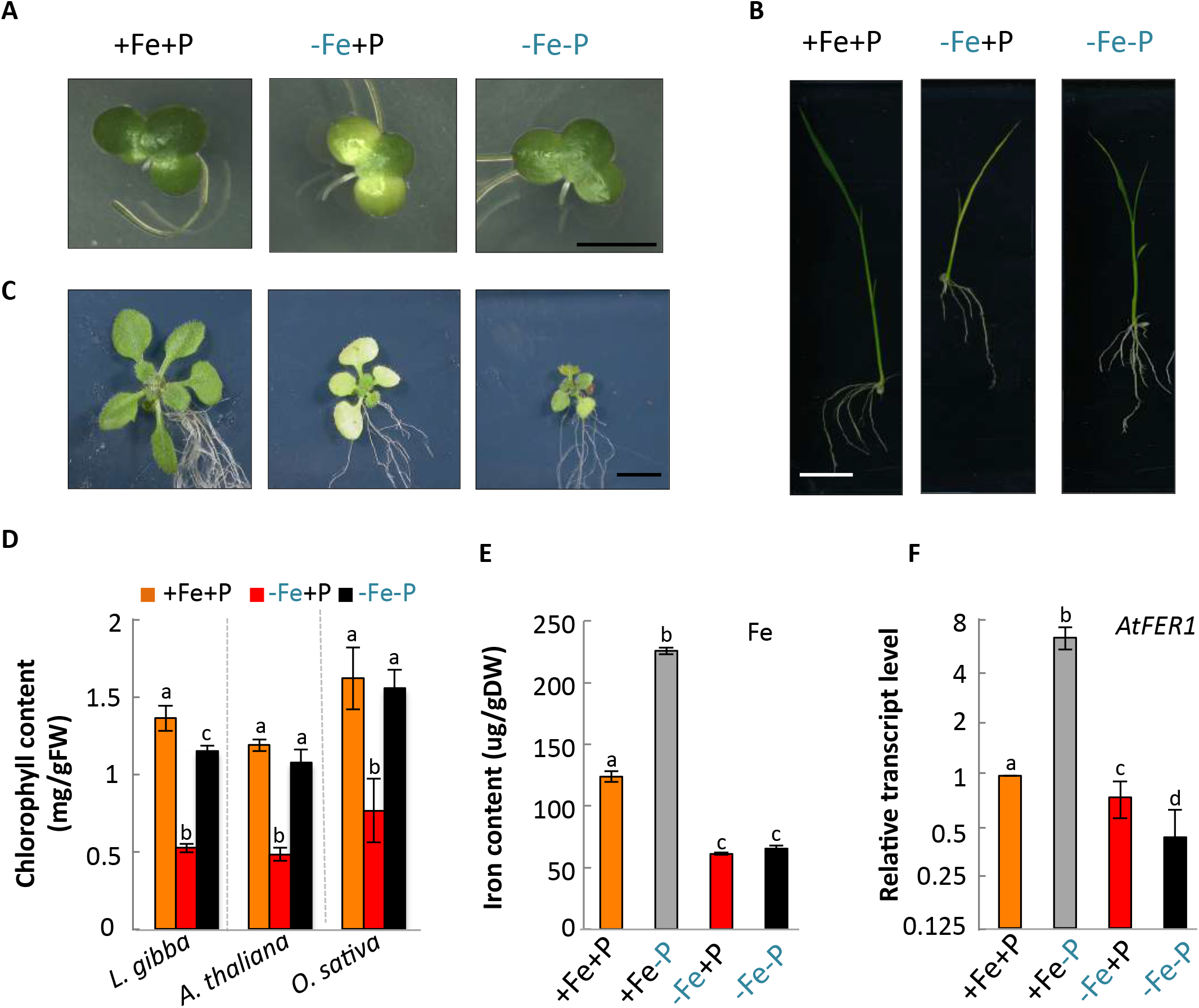
Phosphorus deficiency prevents iron deficiency-induced chlorosis in evolutionarily distant plant species. **A-C**) Duckweed (*Lemna gibba*), rice (*Oryza sativa* cv Nipponbare), and *Arabidopsis thaliana* plants grown on media containing iron and phosphorus (+Fe+P), deficient in iron (-Fe+P), or deficient in both elements (-Fe-P). Representative images of *L. gibba* propagated for 28 days (A), 24-day old rice (B) and 14-day old *A. thaliana* (Col-0) (C) are shown. **D**) Chlorophyll accumulation in *L. gibba, A. thaliana*, and *O. sativa* grown under +Fe+P, -Fe+P, and - Fe-P conditions. Scale bars, 7 mm (A), 10 mm (B), and 5 mm (C). Data shown are from 3 experiments and 3-10 plants per experiment. Error bars represent 95% confidence interval. FW, fresh weight. **E**) Iron content in shoots of *A. thaliana* plants grown on agar plates containing +Fe+P, -Fe+P, or -Fe-P. Data shown are from 3 experiments and 10 plants per experiment. Error bars represent 95% confidence interval. DW, dry weight. **F)** mRNA abundance of *FERRITIN1* (*AtFER1*) relative to *Ubiquitin 10 (At4g05320)* in *A. thaliana* shoots grown under +Fe+P, -Fe+P, or -Fe-P. Data shown from 3 experiments, with above-ground tissue from 5 plants pooled per experiment for RNA extraction. Error bars represent 95% confidence interval. For D-F, letters above bars represent statistically different means at P < 0.05 (one-way ANOVA with a Duncan post-hoc test). Source data are provided as a Source Data file.

To understand the cause of chlorophyll reduction in response to -Fe, we first explored the timing of -Fe sensing and photosynthetic function response. Since -Fe affects chlorophyll accumulation and photosystem II (PSII) activity^13,14^, we monitored their kinetics over 172 hours (h) (Figure 2A-B, Figure S1A-C). Arabidopsis plants were first grown on +Fe+P media for one week, and then transferred to +Fe+P, -Fe+P, or -Fe-P conditions. -Fe+P caused a significant decrease in chlorophyll content observable starting at 52 h after the transfer to -Fe+P (Figure 2A). However, transfer to -Fe-P did not affect chlorophyll content, even at 172 h after the transfer (Figure 2A). To determine how photosynthesis was affected, we measured Fv/Fm, which reflects the quantum yield of photochemistry and is a measure of PSII activity^13,14^. Plants under -Fe+P decreased Fv/Fm observable starting at 52h, indicative of compromised electron transport through PSII, and which coincides with the decrease of chlorophyll accumulation (Figure 2B). By 172 h, PSII activity was substantially reduced under -Fe+P compared to +Fe+P. However, plants under -Fe-P showed slightly lower but stabilized Fv/Fm compared to those in +Fe+P (Figure 2B). These physiological characterizations showed that chlorophyll accumulation and PSII activity were affected by -Fe, and both responses were P-dependent (Figure 2A-B).

**Figure 2.**
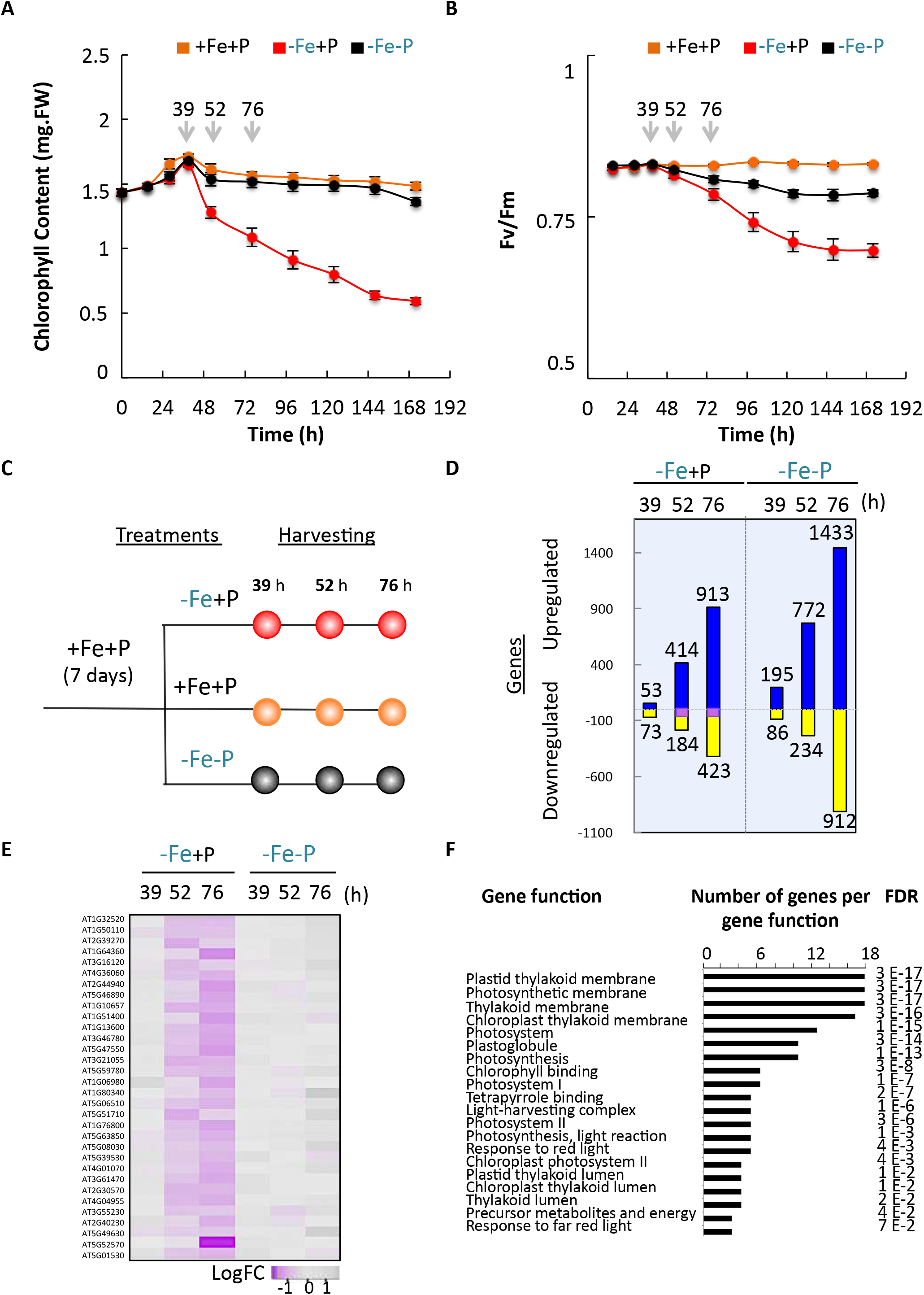
Kinetics of chlorophyll accumulation, photosystem II activity, and transcriptome change in response to iron and phosphorus availability. **A-B)** Chlorophyll content and PSII activity (Fv/Fm) in response to iron and/or phosphate deficiency in *A. thaliana.* Seedlings were grown for 7 days in the presence of iron and phosphorus (+Fe+P) and transferred to three different media: +Fe+P, -Fe+P, or -Fe-P for 15h, 28h, 39h, 52h, 76h, 100h, 124h, 148h, 172h. **A)** Chlorophyll data shown are from 3 experiments, each experiment with 8 plants. Error bars represent 95% confidence interval. **B)** Plants were dark-adapted for 30 min before measuring fluorescence kinetics from a leaf. Fv/Fm data shown are from 3 experiments, each experiment with 16 plants. Error bars represent 95% confidence interval. **C**) Experimental design for transcriptomic studies on *A. thaliana* (Col-0) shoots. Plants were grown in media containing iron and phosphorus (+Fe+P) for 7 days, transferred to three different media (+Fe+P, -Fe+P, or -Fe-P) for 39h, 52h, or 76h, and shoots were harvested for RNA extraction and sequencing. **D**) Global expression analysis of genes in response to -Fe+P and -Fe-P relative to +Fe+P. Numbers of genes displaying at least 2-fold change (p-value < 0.05) in their expression are shown for each condition. The 32 genes that were decreased specifically in -Fe+P but not in - Fe-P relative to +Fe+P at 52h and 76h (highlighted in purple) were used to perform genome-wide association studies. **E**) A heatmap showing gene expression patterns of the 32 genes in -Fe+P and -Fe-P relative to control (+Fe+P) at 39h, 52h, 76h after the transfer. LogFC, log2 fold change. **F**) Gene Ontology analysis for the biological processes (GO-BP) for the 32 genes whose mRNA abundance was specifically decreased by -Fe+P. FDR = false discovery rate. Source data are provided as a Source Data file.

Based on these findings, we selected three time points at 39, 52, and 76 h after the transfer of plants to +Fe+P, -Fe+P, or -Fe-P to conduct a global gene expression analysis in shoots (Figure 2C, Figure S2A-F). We identified genes whose expression levels were either increased or decreased by -Fe+P relative to +Fe+P by at least 2-fold at a p-value < 0.05 (Figures 2D, Table S1). Even more genes were either upregulated or downregulated in -Fe-P conditions relative to +Fe+P (Figures 2D, Figure S3A-C). A total of 671 and 2434 transcripts were uniquely differentially regulated in response to -Fe+P or -Fe-P, respectively, supporting the existence of different signaling pathways under the two conditions (Figure S 3A-C). To identify functions enriched in genes that were differently regulated by -Fe+P or -Fe-P, we performed Gene Ontology (GO) enrichment analysis. Genes specifically downregulated by -Fe-P but not by -Fe+P at 52 h and 76 h (52 genes; Figure S4A, Table S1) showed an enrichment for ribosomal genes (Figure S4B) while upregulated genes (162 genes, Figure S4C, Table S1) revealed an enrichment for genes involved in cation transport, response to water, and ester hydrolysis (Figure S4D, Table S1). On the other hand, GO analysis of the 32 genes specifically downregulated by -Fe+P but not affected by -Fe-P at 52 h and 76 h (Figure 2E, Table S1) revealed an enrichment of genes related to the chloroplast and photosynthesis-related processes (Figure 2F), while upregulated genes (Figure S4E) were enriched for genes related to cellular respiration, oxidation-reduction process, and energy metabolism (35 genes; Figure S4F). Altogether, the transcriptomics analysis indicated that the control of chloroplast function is an integral component of the nuclear transcriptomic response to -Fe, which is dependent on P availability. We also learned that the photosynthesis-related phenotypes we observed under -Fe+P, but not under -Fe-P, could be due to downregulation of key photosynthesis regulators.

To decode the signaling pathways that control expression of the photosynthesis genes in response to -Fe+P, we exploited natural variation in expression of the 32 genes that were down-regulated by Fe deficiency in a P-dependent manner in a worldwide collection of *A. thaliana* accessions ^15^. One way to identify regulatory mechanisms could be to perform expression genome-wide association studies (eGWAS) using the expression level of individual genes across Arabidopsis accessions. Strikingly, we found that expression levels of the 32 genes are predominantly positively associated with each other across 727 Arabidopsis accessions (Figure S5A) ^15^. We then performed Principal Component Analysis (PCA) to reduce the dimensionality of these expression data. PC1 explained 89.5% of the variation in expression of these genes across Arabidopsis accessions (Figure S5B). The contribution of each accession to PC1 was then used to perform a genome-wide association study (GWAS) (Figure 3A). Our GWAS analysis detected 38 QTLs containing 145 candidate genes, based on a 20-kb window per QTL and using a 5% false discovery rate (FDR) threshold, with the highest peak (Chromosome 2) occurring in an intergenic region (Figure 3A). In this study, we followed up one of the QTLs that contained the inorganic phosphate transporter *PHT4;4* (AT4G00370) (Figure 3A, Figures S6 and S7) given its role in ascorbic acid (AsA) transport into chloroplasts, which was proposed to be important for maintaining the xanthophyll cycle for dissipation of excessive light energy to heat in photosynthesis ^16^ To determine if PHT4;4 has any role in -Fe+P dependent chlorosis, we examined mutants with a null *pht4;4* allele. Under -Fe+P, chlorophyll was significantly reduced in *pht4;4* mutants, similar to wild type plants (Figure 3B-C). However, -Fe-P conditions failed to recover chlorosis and chlorophyll reduction in *pht4;4* mutants, unlike wild type plants (Figure 3B-C). Introduction of the wild type *PHT4;4* allele into a *pht4;4* mutant background complemented these phenotypes (Figure 3B-C). The chlorotic phenotype of the *pht4;4* mutant under -Fe-P suggested that transport of AsA into chloroplasts could be important for the “stay green” phenotype.

**Figure 3.**
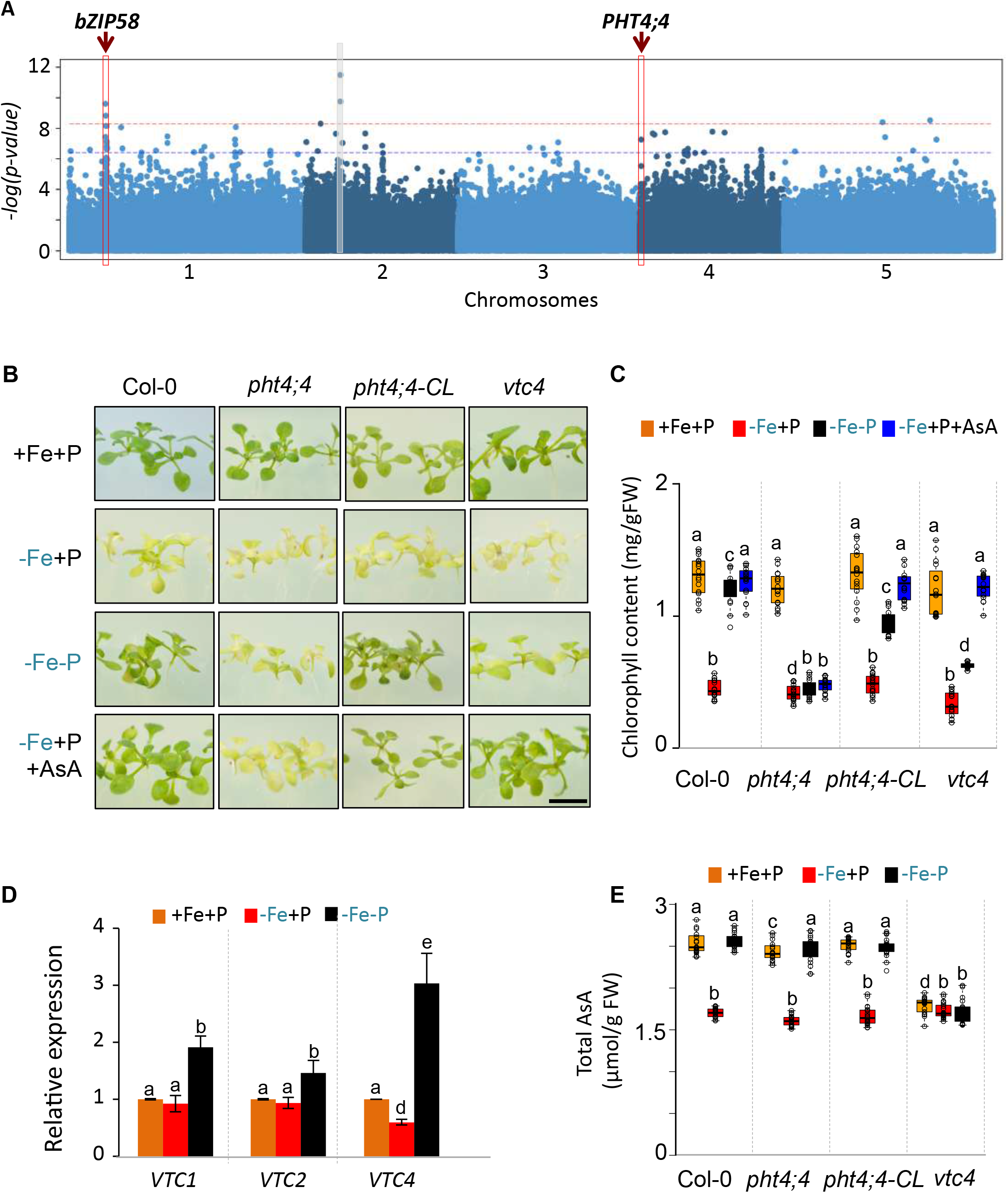
PHT4;4 prevents chlorosis under the combined deficiency of iron and phosphorus. **A)** A Manhattan plot for genome-wide association mapping using principal component 1 that explained 89.5% of expression variation of the 32 photosynthesis related genes across 727 *A. thaliana* accessions ^15^. The five chromosomes are depicted by light and dark blue colors. Dashed lines correspond to FDR 5% threshold (blue) and Bonferroni α = 0.05 (red). The light grey rectangle highlights a significant association located in an intergenic region (SNP: 4493712). Two significant associations that were followed up in this study are highlighted in red rectangles. **B**) Representative images of wild-type Col-0, *pht4;4, vtc4*, and a line expressing genomic *PHT4;4* in *pht4;4* (*pht4;4-CL*) grown for 7 days in the presence of iron and phosphorus (+Fe+P) and transferred to three different media: +Fe+P, - Fe+P, or -Fe-P for 7 additional days. Scale bars: 7 mm **C**) Total chlorophyll content in Col-0, *pht4;4*, *vtc4*, and *PHT4;4-CL* grown under -Fe+P, +Fe-P, -Fe-P, or - Fe+P+AsA. Data shown are from 10 plants conducted in three independent experiments. **D**) mRNA abundance of *VTC* genes (*VTC1, VTC2*, and *VTC4)* relative to *Ubiquitin 10 (At4g05320)* in shoots of *A. thaliana* Col-0 plants grown in the presence of iron and phosphorus (+Fe+P) for 7 days and transferred to +Fe+P, - Fe+P, -Fe-P or -Fe-P+AsA for 52h. Data shown are from 3 experiments. Error bars represent 95% confidence interval. **E**) Total ascorbic acid (AsA) content in Col-0, *pht4;4*, *vtc4*, and *PHT4;4-CL* plants grown for 7 days in the presence of iron and phosphorus (+Fe+P) and transferred to +Fe+P, -Fe+P, or -Fe-P for 52h. Data shown are from 3 experiments, each with 16 plants. In box plots (C, E) center lines show sample medians; box limits indicate the 25th and 75th percentiles; whiskers extend 1.5 times the interquartile range from the 25th and 75th percentiles. For C-E, letters above bars or boxes represent statistically different means at P < 0.05 (one-way ANOVA with a Duncan post-hoc test). Data points are plotted as open circles. Source data are provided as a Source Data file.

To test our hypothesis about the role of AsA in preventing chlorosis under -Fe-P conditions, we first assessed how Fe and P availabilities regulate the expression of *VITAMIN C (VTC)* enzymes involved in AsA biosynthesis in plants ^17^. Our RNA-seq analysis identified that -Fe-P caused a 2 to 3-fold increase in *VTC1, VTC2*, and *VTC4* expression, which we confirmed using qRT-PCR (Figure 3D). However, -Fe in the presence of P (-Fe+P) caused a 2-fold decrease in the mRNA abundance of *VTC4* (Figure 3D). VTC4 is the final enzyme in the AsA biosynthesis pathway ^17^. This prompted us to test the effect of the absence of *VTC4* on chlorophyll accumulation under -Fe+P and -Fe-P conditions. Under -Fe-P, mutants with a *vtc4* null allele were still chlorotic, similarly to *pht4;4* and in contrast to wild type plants (Figure 3B-C). These data show that AsA contributes to preventing chlorosis in -Fe-P conditions.

Next, we tested whether the chlorotic phenotype is due to variations in AsA levels. In wild type, AsA levels decreased significantly under -Fe+P at 52 h after the transfer relative to control (+Fe+P), whereas no change was detected under -Fe-P (Figure 3E), suggesting that AsA levels were associated with -Fe-mediated chlorosis. To test whether AsA levels were associated with chlorosis in general, we measured AsA levels in AsA synthesis (*vtc4*) mutant plants. Under +Fe+P, *vtc4* plants accumulated 35% less AsA than wild type plants, and AsA levels remained unchanged in response to -Fe+P or -Fe-P stress (Figure 3E). However, *vtc4* plants did not show the chlorotic phenotype under +Fe+P, which indicated that the level of AsA contributed to the chlorotic phenotype specifically under -Fe and this contribution was dependent on P availability. In addition, the AsA transporter (*pht4;4*) mutants showed similar AsA levels as the wild type even though *pht4;4* plants were still chlorotic in - Fe-P (Figure 3E). To determine whether AsA accumulation in the cell or its transport to the chloroplast is associated with the development of chlorotic phenotype in - Fe+P, we tested the effect of an exogenous supply of AsA in wild type, *vtc4*, and *pht4;4* plants (Figure 3B-C). Exogenous AsA alleviated the chlorosis caused by - Fe+P in wild type and *vtc4* mutant plants. However, *pht4;4* mutants failed to stay green under -Fe+P+AsA conditions (Figure 3B-C), indicating that the transport of AsA to the chloroplast is required for -P mediated ‘stay green’ phenotype under Fe deficiency. Our results showed that -P prevents the downregulation of *VTC4* by -Fe and associated changes in AsA accumulation, and that the PHT4;4-mediated transport of AsA to chloroplasts is required for the maintenance of chlorophyll content under combined deficiency of Fe and P.

We next asked whether *PHT4;4-mediated* AsA transport to the chloroplast is important for regulation of the photosynthesis-related genes that were specifically down-regulated by -Fe in a P-dependent manner. First, we tested the effects of *PHT4;4* inactivation on the expression of these photosynthesis related genes using qRT-PCR (Figure 4A). While -Fe+P significantly downregulated the mRNA abundance of these genes in wild type plants (Col-0), -Fe-P prevented this response (Figure 4A). Furthermore, adding AsA to -Fe+P mimicked -Fe-P response in preventing down-regulation of the photosynthesis genes. Under -Fe+P, *pht4;4* mutant plants showed a decrease in the mRNA abundance of these genes comparable to wild type plants (Figure 4A). However, in contrast to the wild type, these genes were still downregulated in *pht4;4* plants under -Fe-P as well as -Fe+P supplemented with AsA (Figure 4A). Taken together, these data indicate that the transport of AsA to chloroplasts via PHT4;4 is central to preventing the downregulation of photosynthesis-related genes under -Fe-P.

**Figure 4.**
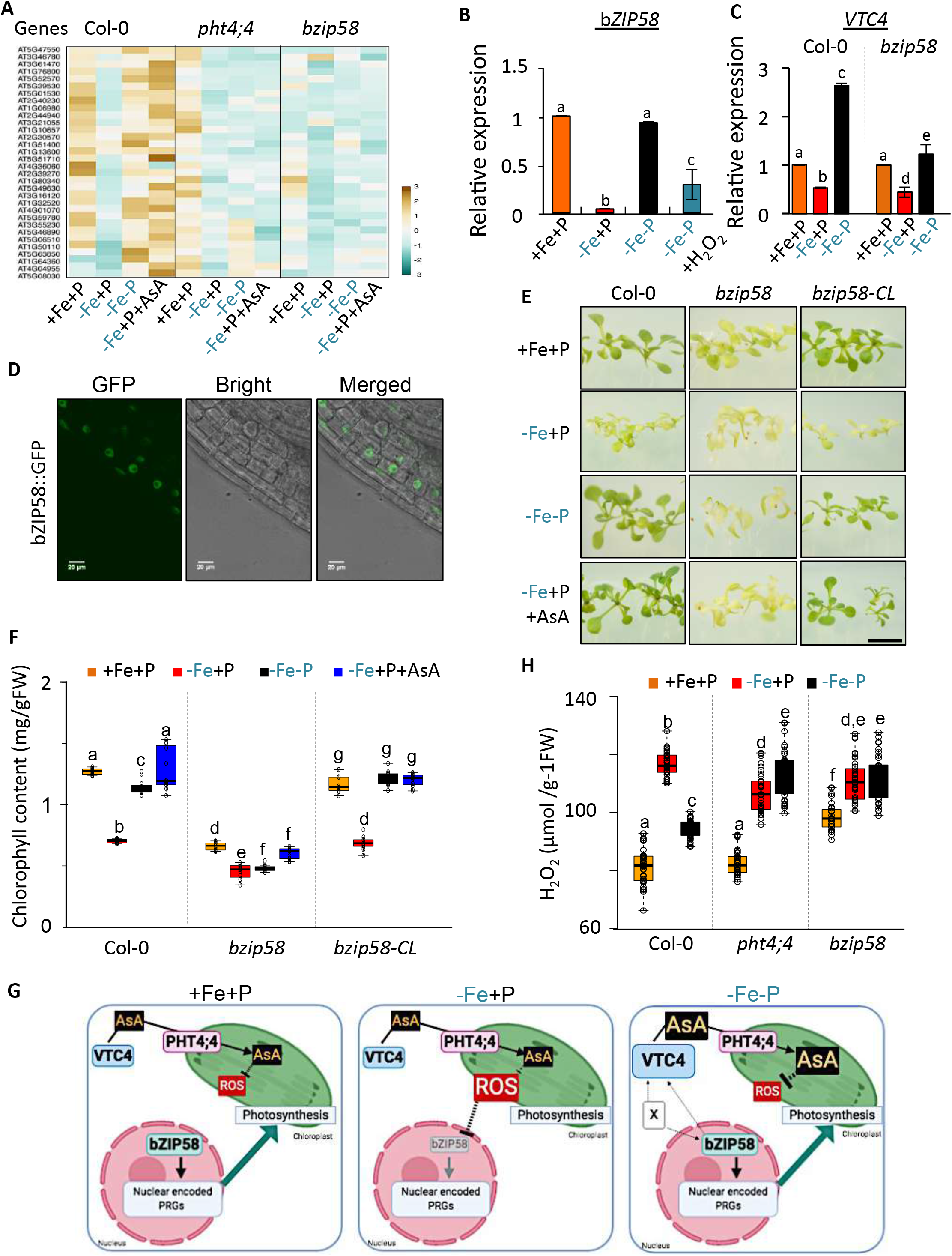
bZIP58 regulates photosynthesis related genes and chlorophyll accumulation. **A)** mRNA abundance of 32 photosynthesis-related genes relative to *Ubiquitin 10* in shoots of *A. thaliana* Col-0, *pht4;4*, and *bzip58* plants grown for 7 days in the presence of iron and phosphorus (+Fe+P) and transferred to +Fe+P, -Fe+P, - Fe-P, or -Fe-P+AsA for 52h. Data were averaged from three independent experiments. Scale bar represents the relative mRNA abundance. **B)** mRNA abundance of *bZIP58* relative to *Ubiquitin* 10 in *A. thaliana* Col-0 shoots of plants grown for 7 days on +Fe+P and transferred to +Fe+P, -Fe+P, -Fe-P, or +Fe+P+H_2_O_2_ for 52h. Data are shown from 3 experiments. Error bars represent 95% confidence interval. **C**) *VTC4* mRNA abundance relative to *Ubiquitin* 10 in the shoots of Col-0 and *bzip58* mutants grown in the presence of +Fe+P for 7 days and transferred to +Fe+P, -Fe+P, or -Fe-P for 52h. Data shown from 3 experiments. Error bars represent 95% confidence interval. **D)** Confocal microscopy images of p35S::bZIP58::GFP expressing plants (scale bar = 20*μ*m) grown for 7 days under +Fe+P. **E**) Representative images of Col-0, *bzip58*, and a line expressing genomic *bZIP58* in *bzip58* mutants (*bzip58-CL*) grown for 7 days in +Fe+P and transferred to +Fe+P, -Fe+P, or -Fe-P for 7 additional days. Scale bars: 7 mm. **F**) Total chlorophyll content in Col-0, *bzip58*, and *bZIP58-CL* plants grown for 7 days in +Fe+P and transferred to -Fe+P, +Fe-P, -Fe-P, or -Fe-P+AsA for 7 days. Data shown are from 4 experiments. **G**) A schematic model delineating a signaling pathway that integrates Fe and P availability cues to regulate chlorophyll accumulation and photosynthesis genes. Fe deficiency (-Fe+P) causes a decrease in the expression of *bZIP58* that is central to controlling the transcription of nuclear-encoded photosynthetic genes. P limitation under Fe deficiency (-Fe-P) prevents this downregulation of *bZIP58* and induces *VTC4*. The induction of *VTC4* expression requires bZIP58, whose effect could be direct or indirect, represented here by ‘X’. We propose that induction of *VTC4* increases ascorbic acid in the chloroplast mediated by PHT4;4. We hypothesize that the increase of ascorbic acid level prevents ROS accumulation, thus maintaining the expression of bZIP58 and its downstream photosynthesis genes leading to the ‘stay green’ phenotype. **H**) Accumulation of H_2_O_2_ (a type of ROS) in shoots of Col-0, *bzip58*, and *bZIP58-CL* plants grown for 7 days in +Fe+P and transferred to +Fe+P, -Fe+P, or -Fe-P for 52h. Data shown from 12 experiments. In box plots (F, H) center lines show sample medians; box limits indicate the 25th and 75th percentiles; whiskers extend 1.5 times the interquartile range from the 25th and 75th percentiles. For B, C, F, H, letters above bars or boxes represent statistically different means at P < 0.05 (one-way ANOVA with a Duncan post-hoc test). Data points are plotted as open circles. Source data are provided as a Source Data file.

How does -Fe+P affect the expression of photosynthesis-related genes? To look for potential transcriptional regulators of these genes, we examined the candidate genes from the GWAS analysis. We found that *bZIP58* (AT1G13600) (Figure 3A, Figure S6-S7), a putative transcription factor, underlies one of the strongest QTL peaks (Figure 3A). To test whether *bZIP58* responds to Fe and P availability, we performed qRT-PCR of *bZIP58* under various Fe and P availability. *bZIP58* was strongly downregulated by -Fe, and this down-regulation depended on P availability (Figure 4B). In addition, bZIP58 was partially required to induce *VTC4* expression under -Fe-P conditions (Figure 4C). This led us to examine the contribution of bZIP58 in regulating the -Fe+P specific photosynthesis-related genes under +Fe+P, -Fe+P, -Fe-P conditions (Figure 4A). Mutants with the *bzip58* null allele showed a remarkable constitutive decrease in the expression of these 32 photosynthesis-related genes (Figure 4A). bZIP58 localizes to the nucleus (Figure 4D), which is consistent with a role as a transcription factor. Taken together, these findings support the idea that bZIP58 is a key regulator of photosynthesis-related genes, and its absence could alter chlorophyll accumulation regardless of Fe and P availability. Genetic inactivation of *bZIP58* indeed causes a constitutive decrease in chlorophyll content, and the mutant line is chlorotic (Figure 4E-F). The expression of *bZIP58 gene* in *bzip58* plants complements the constitutive chlorosis phenotype, and the complemented line responds to Fe and P availability similarly to wild type plants (Figure 4E-F). Furthermore, AsA supplementation could not rescue the chlorotic phenotype of *bzip58* mutants (Figure 4E-F), indicating that pZIP58 lies downstream of AsA action. These data show that bZIP58 controls the expression of photosynthesis-related genes and is transcriptionally regulated in response to -Fe depending on P availability, likely by mediating the perception of AsA.

Based on our findings, we propose a model to explain how P availability modulates Fe deficiency-induced chlorosis (Figure 4G). -Fe+P causes a decrease in the expression of AsA biosynthesis gene *VTC4* (Figure 3D). Exogenous AsA supply prevents the development of chlorosis in *vtc4* under -Fe-P, but not in the *pht4;4* that transports AsA to the chloroplast (Figure 3B). We thus propose that AsA deficiency in the chloroplast under Fe limited conditions triggers chlorosis. How does AsA in the chloroplast affect photosynthesis? AsA has an antioxidizing action that detoxifies reactive oxygen species (ROS) through its scavenging properties ^18^, thus making ROS a potential signaling molecule^19-21^ capable of modulating the expression of photosynthesis-related genes through bZIP58. To test this hypothesis, we measured the relative amount of ROS accumulation in shoots of wild type, *pht4;4* and *bzip58* plants under various Fe and P availability. -Fe+P caused a 2-fold increase in ROS accumulation in shoots of wild type plants, which partially depended on P availability (Figure 4H). *pht4;4* plants displayed comparable ROS accumulation to that of the wild type under +Fe+P and -Fe+P. However, *pht4;4* plants accumulated significantly higher ROS than wild type plants under -Fe-P (Figure 4H). In addition, *bzip58* mutant plants displayed a constitutive increase in ROS accumulation (Figure 4H). To check whether ROS in turn can regulate the expression of *bZIP58*, we quantified *bZIP58* expression in response to foliar application of H_2_O_2_. ROS treatment caused a 4-fold decrease in *bZIP58* transcript accumulation (Figure 4B). Collectively, our results support the idea that under simultaneous Fe and P deficiency, AsA accumulation in the chloroplast prevents chlorosis by modulating ROS levels that may control the expression of photosynthesis genes via a putative transcription factor bZIP58 (Figure 4G). How ROS acts as a retrograde signal to alter nuclear gene expression to control photosynthesis under Fe and P limitation remains to be determined, though we now have several molecular targets with which to explore this field. Modulation of the discovered pathway could have a direct impact on plant growth in the field by improving plant photosynthetic activity while reducing nutrient supply.

## Supplemental Figures

**Figure S1.**
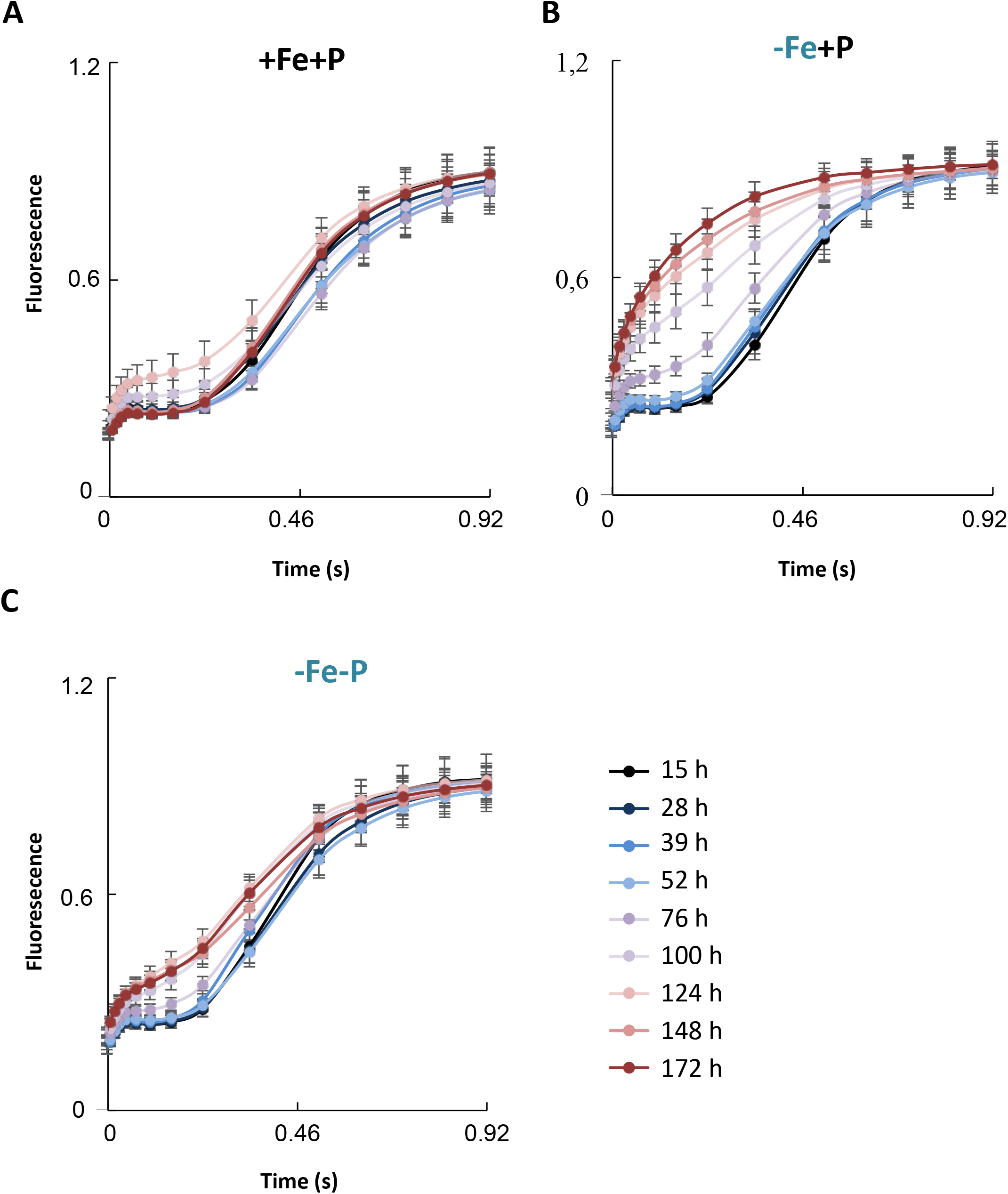
Photosystem II activity in response to Fe and/or P deficiency in *A. thaliana*. Seedlings were grown for 7 days in the presence of Fe and P (+Fe+P) and transferred to three different media: +Fe+P, -Fe+P, or -Fe-P for 15h, 28h, 39h, 52h, 76h, 100h, 124h, 148h, 172h. Plants were dark-adapted for 30 min before measuring the kinetics of fluorescence from a leaf. Data shown are from 3 experiments and 13 to 16 plants were measured per experiment. Error bars represent 95% confidence interval. Source data are provided as a Source Data file.

**Figure S2.**
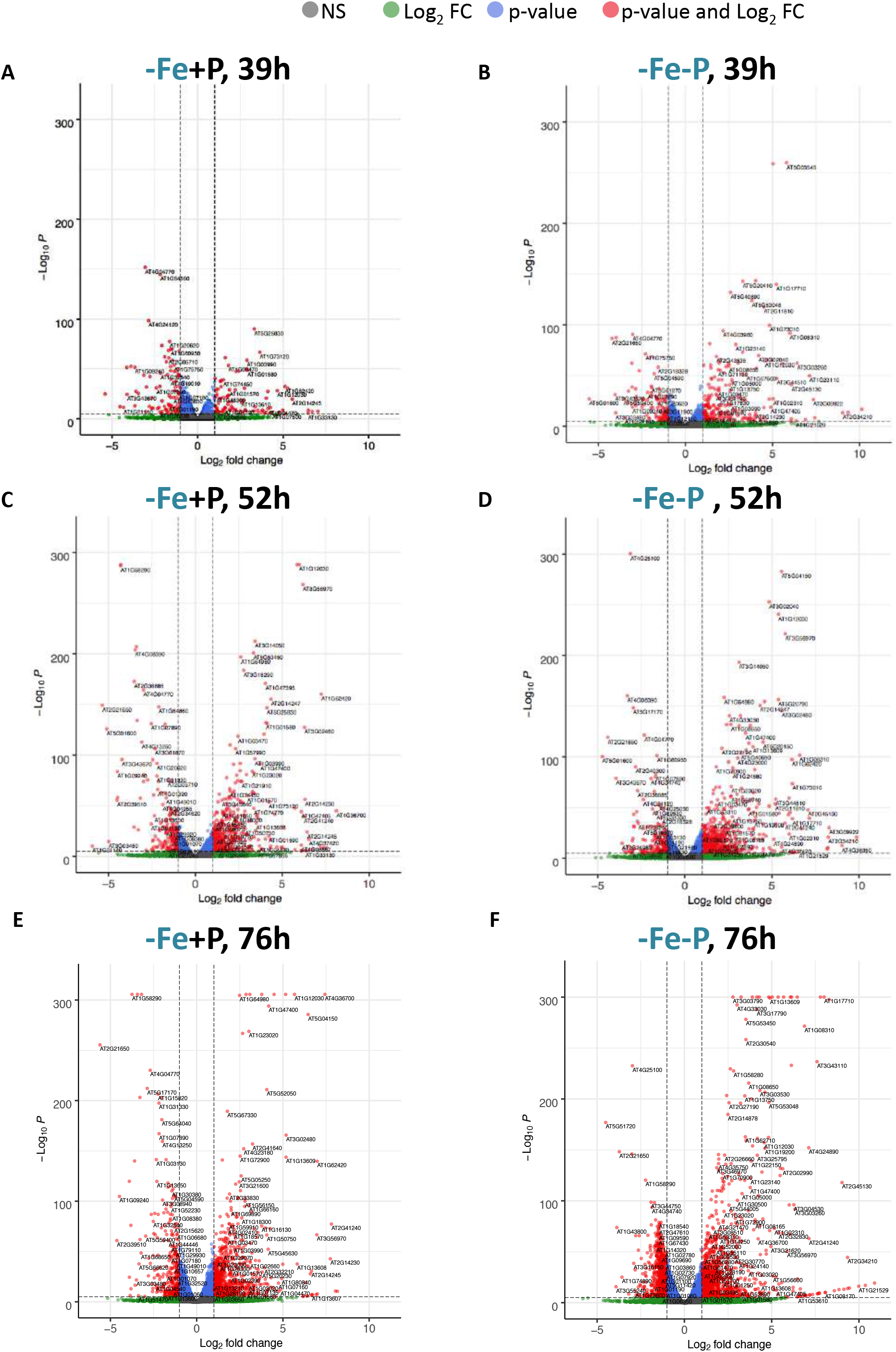
Transcriptome kinetics of *A. thaliana* in response to Fe and/or P deficiency. Volcano plots of individual transcript abundance in wild-type plants (Col-0) grown in -Fe+P (A,C,E) or -Fe-P (B,D,F) relative to +Fe+P. Shoot samples were collected from plants grown for 39h (A,B), 52h (C,D), or 76h (E,F). x-axis: foldchanges; y-axis: adjusted p-values based on Benjamini-Hochberg correction; Both axes use log scales. Red: log2Fold-Change>|2| and -log10P>6; Blue: log2FoldChange<|2| and -log10P>6; Green: log2FoldChange>|2| and -log10P<6; Grey: log2FoldChange<|2| and -log10P<6.

**Figure S3.**
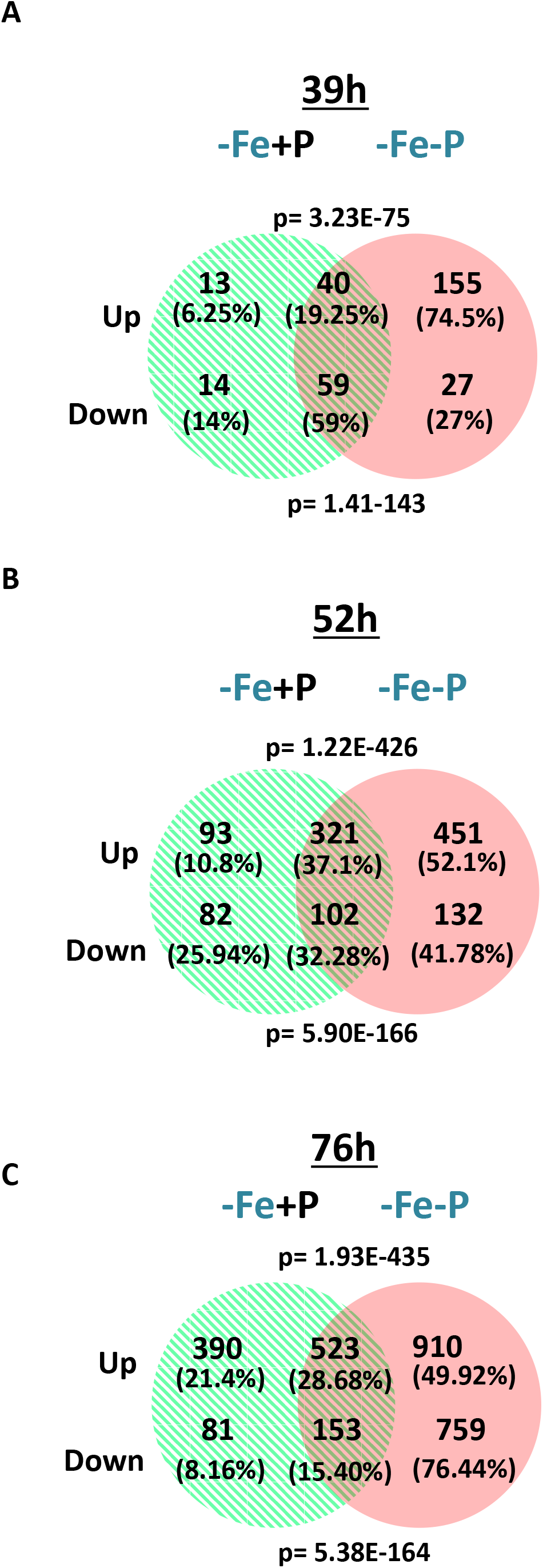
Common and unique genes regulated by Fe and/or P deficiency. **A-C)** Venn diagrams show the genes that are commonly and specifically increased or decreased in abundance in shoots of *A. thaliana* wild type (Col-0) plants grown in +Fe+P for 7 days and transferred to -Fe+P or -Fe-P for 39h, 52h, or 76h relative to those transferred to +Fe+P (fold change >2, p<0.05). The Venn diagram was constructed using a web-based tool (http://bioinformatics.psb.ugent.be/webtools/Venn/). p = p-values from hypergeometric testing.

**Figure S4.**
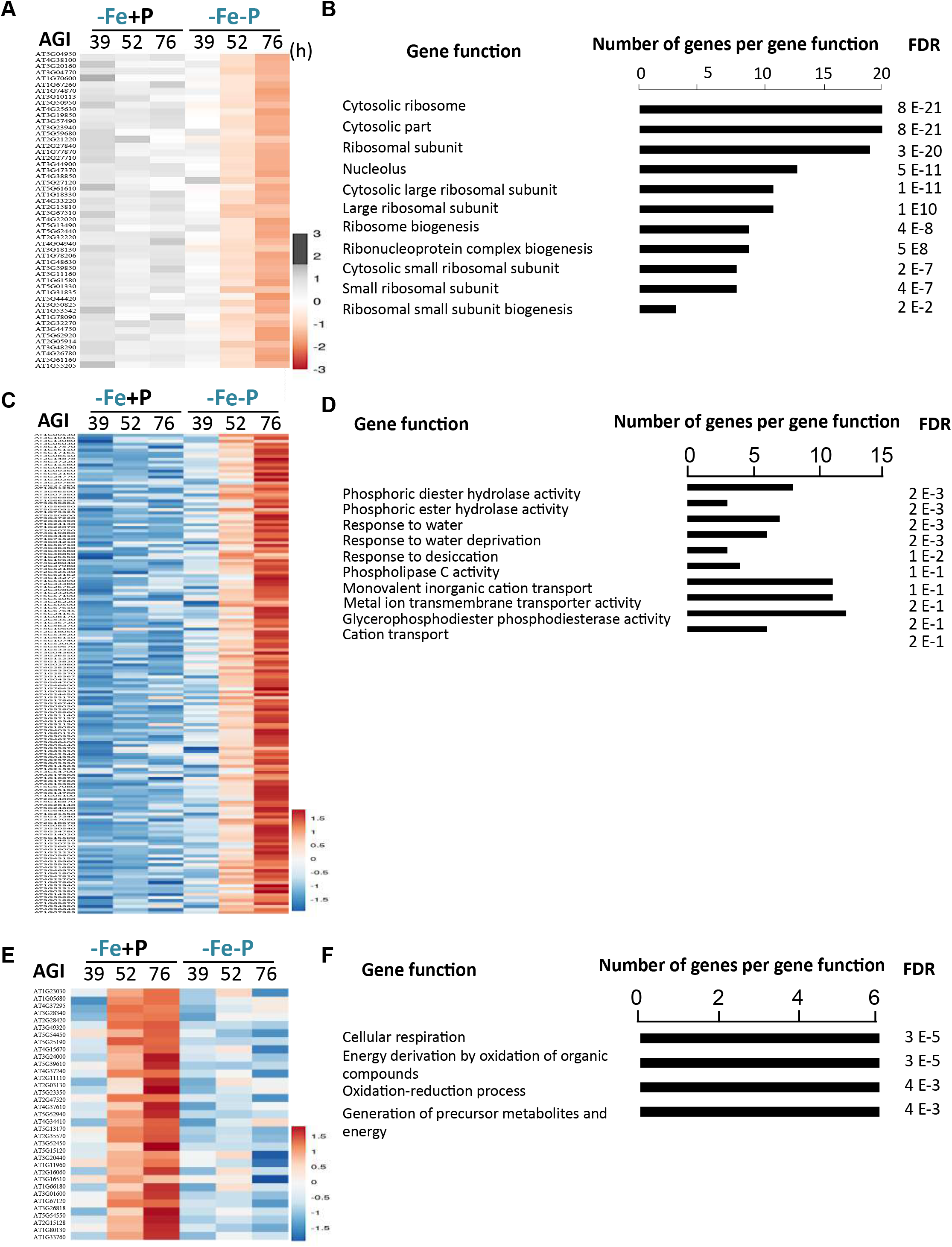
Gene Ontology enrichment analysis of genes specifically regulated by Fe and/or P deficiency. **A, C)** Heatmaps representing changes in the expression of genes that were specifically decreased in abundance under -Fe-P but not -Fe+P relative to +Fe+P (**A**) and increased in abundance in -Fe-P but not in -Fe+P relative to +Fe+P (**C**). **E**) A heatmap representing the expression of genes specifically upregulated by -Fe+P but not by -Fe-P relative to +Fe+P. **B, D, F**) Gene Ontology enrichment for biological processes (GO-BP) in the genes that were specifically decreased by -Fe-P relative to +Fe+P (**B**), specifically increased by -Fe-P relative to +Fe+P (**D**), and specifically increased by -Fe+P relative to +Fe+P (**F**) using GENEMANIA ^22^. Number of genes in each functional category and adjusted p-values for the enrichment are shown. FDR = false discovery rate.

**Figure S5.**
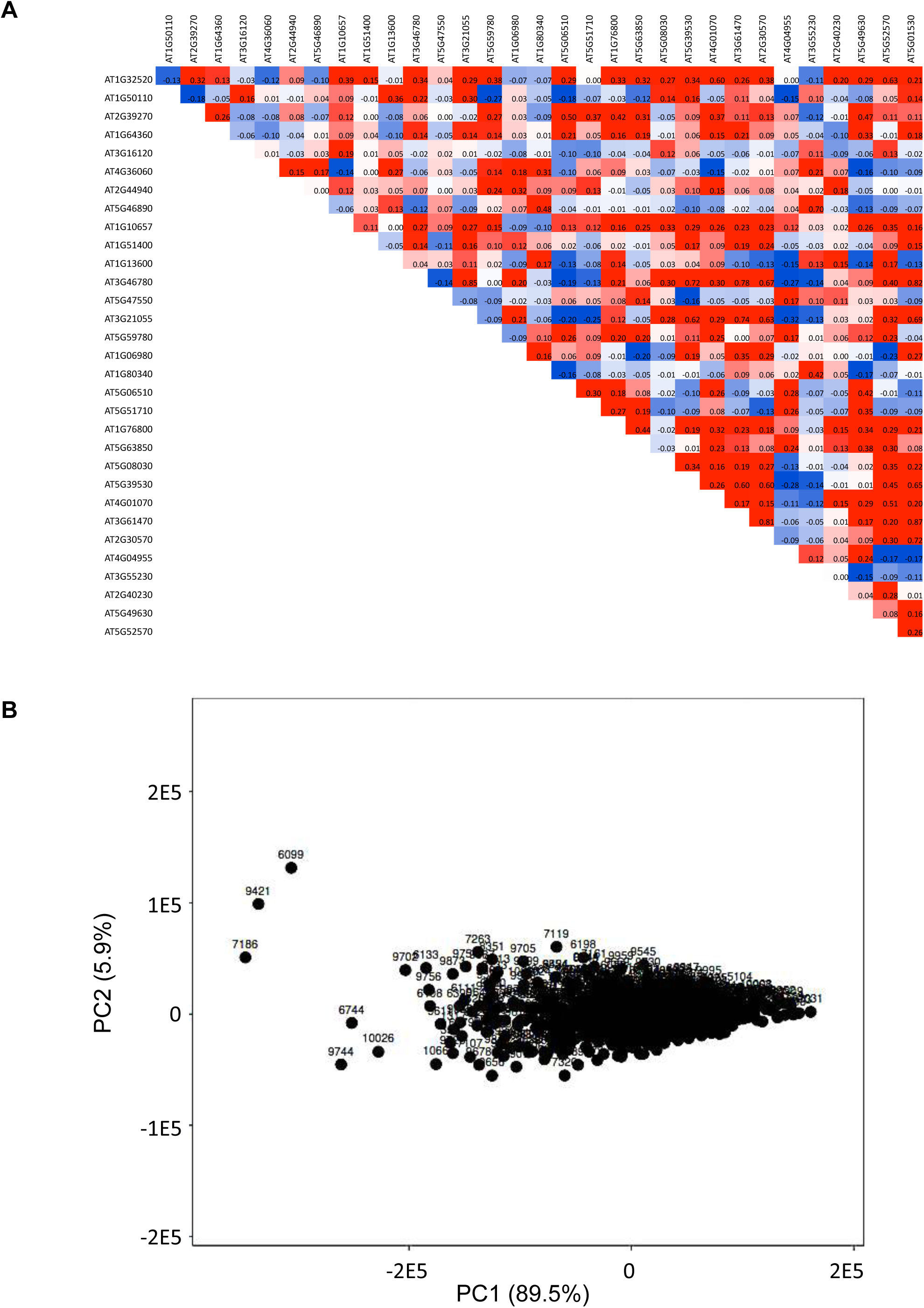
Correlation in the expression of the 32 photosynthesis genes specifically downregulated by -Fe+P across 727 *A. thaliana* accessions. **A**) A heatmap of pairwise correlations (Pearson’s correlation coefficient) in the 32 genes across 727 Arabidopsis accessions grown under control conditions. The correlations were calculated using normalized read counts. **B**) Principal Component (PC) Analysis was performed using the expression of 32 genes in 727 Arabidopsis accessions. X- and Y-axes show PC 1 and PC 2 that explain 89.5% and 5.9 % of the total variance, respectively.

**Figure S6.**
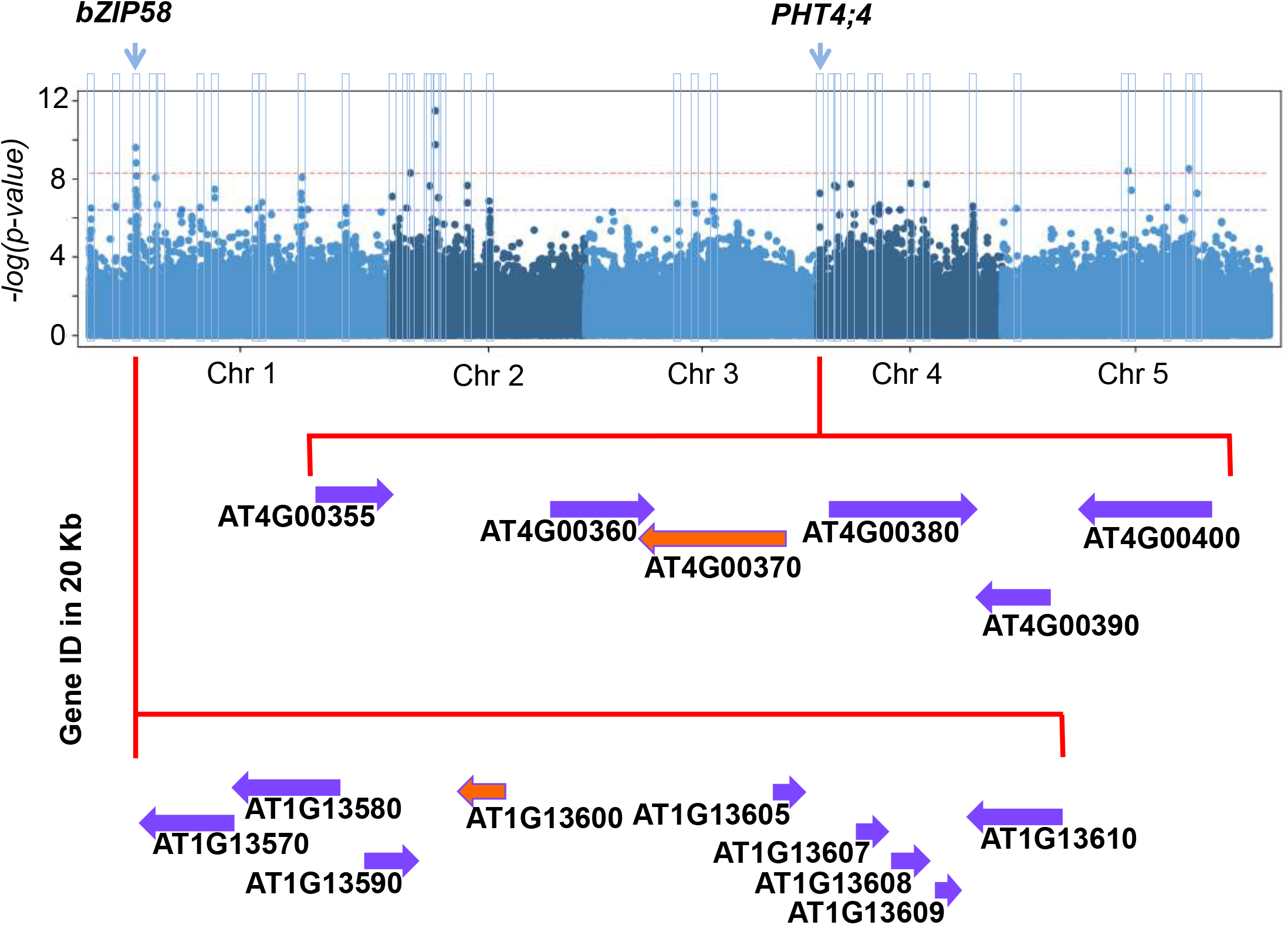
A close-up view of chromosomes 1 and 4 around bZIP58 and PHT4;4 respectively. A Manhattan plot for genome-wide association mapping using PC 1 of the expression profile of the 32 photosynthesis genes across 727 accessions. The five *A. thaliana* chromosomes are depicted by light and dark blue colors. Blue and red horizontal dashed lines correspond to FDR 5% threshold and Bonferroni α = 0.05, respectively. Light blue rectangles indicate the significant SNPs identified in this study. Below the Manhattan plot shows gene models located within a 20-kb genomic region surrounding the two QTLs pursued in this study. Source data are provided as a Source Data file.

**Figure S7.**
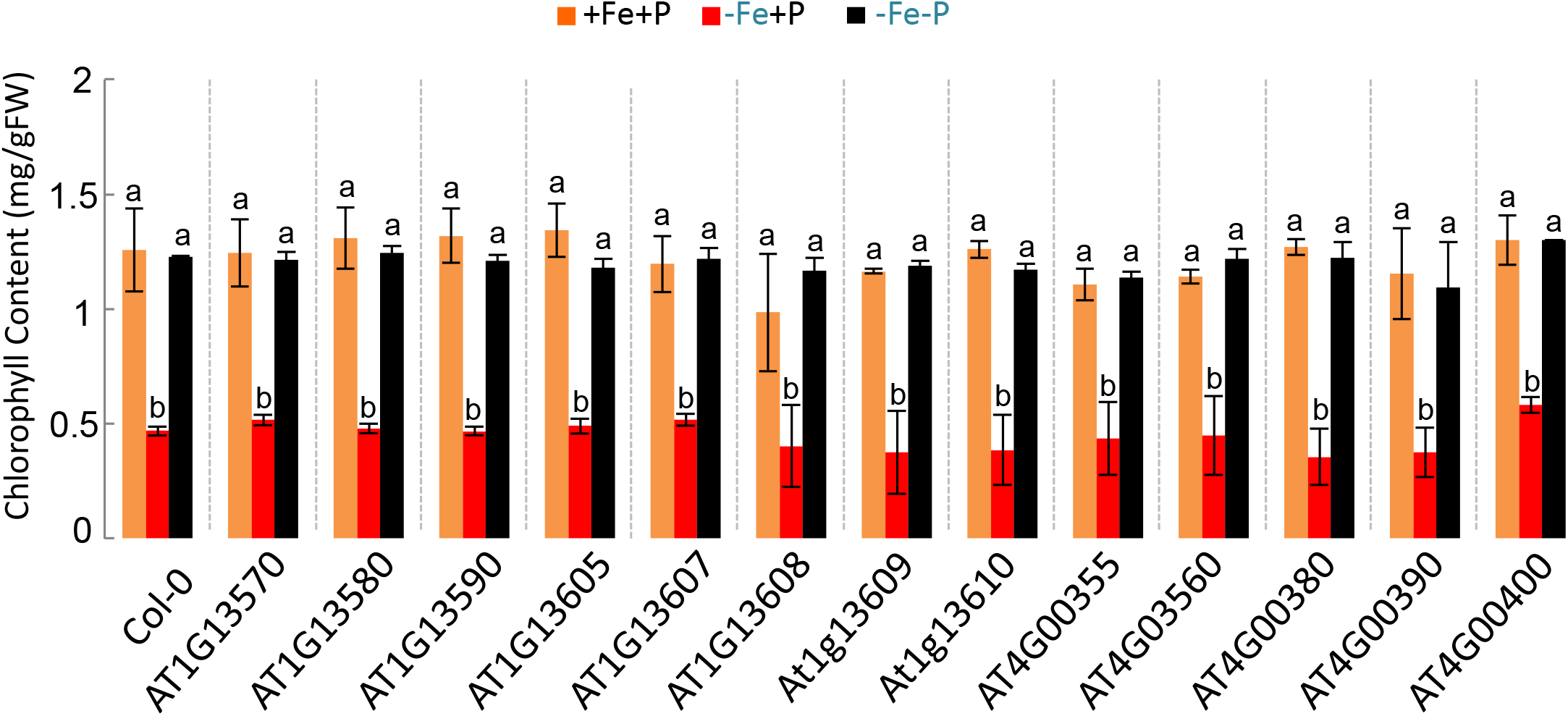
Effects of Fe and/or P availability on chlorophyll content in *A. thaliana* Col-0 and mutants of candidate genes identified using GWAS. Chlorophyll content in Col-0 (CS60000), T-DNA insertion mutant lines in AT1G13570 (SALK_139877), AT1G13580 (SALK_150849), AT1G13590 (SALK_063177), AT1G13605 (SALK_087271), AT1G13607 (SALK_130208), AT1G13608 (SALK_023173), AT1G13609 (SAIL_1243_E04), AT1G13610 (SAIL_897_D11), At4g00355 (N469134), AT4G00360 (SALK_128714), AT4G00380 (SAIL_842_E09), AT4G03585 (SALK_ 128714), At4g00390 (SAIL_313_F07), and At4g00400 (SAIL_633_E10) grown for 7 days in the presence of iron and phosphorus (+Fe+P) and transferred to +Fe+P, -Fe+P, or -Fe-P for an additional week. Data shown from 3 experiments. Error bars represent 95% confidence interval. Letters a and b indicate significantly different values at p <0.05 determined by one-way ANOVA and Tukey’s honest significant difference (HSD) tests. Source data are provided as a Source Data file.

### Supplemental tables

**Table S1. Differentially expressed genes in response to iron and/or phosphorus deficiency in *A. thaliana*.** Gene transcript levels were determined in shoots of Col-0 plants grown in control (+Fe+P) condition for 7 days and then transferred to -Fe+P, -Fe-P, or +Fe+P for 39h, 52h, or 76h.

**Table S2.** List of primers used in this study.

## Materials and Methods

### Plants and growth conditions

Seeds of *Arabidopsis thaliana* wild type (ecotype Columbia, Col-0, CS60000) and knock-out mutant lines SALK_139877 (AT1G13570), SALK_150849 (AT1G13580), SALK_063177 (AT1G13590), N571881 (AT1G13600), SALK_087271 (AT1G13605), SALK_130208 (AT1G13607), SALK_023173 (AT1G13608), SAIL_1243_E04 (AT1G13609), SAIL_897_D11 (AT1G13610), N469134 (At4g00355), SALK_128714 (AT4G00360), N469134 (AT4G00370), SAIL_842_E09 (AT4G00380), N866595 (At4g00390), SAIL_633_E10 (At4g00400) and SALK_077222 (AT3G02870) were obtained from the Nottingham Arabidopsis Stock Centre (NASC). Homozygous mutant lines were confirmed by PCR using the primers listed in Table S2. bZIP58 complemented lines (bZIP58-CL) were generated by expressing 3896 bp genomic DNA containing *bZIP58* in the *bzip58* mutant background (NASC, N571881). Complementation of *pht4;4* mutant plants (PHT4;4-CL) was obtained by expressing 6450 bp genomic DNA containing *PHT4;4* in the *pht4;4* mutant background (NASC, N469134). Arabidopsis plants were grown on control (+Fe+P) plates containing 1.249 mM KH_2_PO_4_; 0.25 mM Ca(NO_3_)_2_; 0.5 mM KNO_3_; 1 mM MgSO_4_; 100 μM FeSO_4_.7H_2_O; 30 μM H_3_BO_3_; 1 μM ZnCl_2_; 10 μM MnCl_2_; 1 μM CuCl_2_; 0.1 μM (NH_4_)_6_Mo_7_O_24_; and 50 μM KCl; 0.05% 2-(N-morpholino)ethanesulfonic acid (MES), without sucrose supplementation, and 0.8% washed agar. The agar was washed 3 times with 50 mM EDTA, pH 5.7, with continuous stirring for 16 h, then washed 6 times with Milli-Q de-ionized water for 2 hours to reduce mineral contamination ^23^. P-deficient media contained 12.49 μM KH_2_PO_4_ (+Fe-P). Fe-free media was obtained by omitting FeSO_4_.7H_2_O from the growth media (-Fe+P). P- and Fe-deficient media contained 12.49 μM KH_2_PO_4_ (+Fe-P), and no FeSO_4_.7H_2_O (-Fe-P). Seeds were stratified at 4°C for 3 days and grown on vertical agar plates in a growth chamber with 22 °C, 24 h of light at 100 μmol m^-2^s^-1^ fluorescent illumination. *Lemna gibba* (duckweed) plants used during this study were obtained from Duckweeds stock center (stock number 29-DWC131) at Rutgers University (USA). Duckweed plants were grown in 1X Schenk & Hildebrandt (SH) hydroponic medium containing 0.05% 2-(N-morpholino)ethanesulfonic acid (MES) and 1% sucrose, and pH adjusted to 5.7. For experiments with duckweeds, P-deficient and Fe-deficient media contained 1% NH_4_H_2_PO_4_ and 1% FeSO_4_7H_2_O, respectively, of 1X SH media. Media were changed every 7 days. The growth condition was 22 °C and 24 h of light at 80 μmol m^-2^s^-1^. Rice (*Oryza sativa* cv Nipponbare) plants were grown hydroponically in 0.25X Yoshida media^24^ under light/dark cycle of 14/10 h, and temperature of 28/25 °C. Single (-P or -Fe) and combined (-P-Fe) nutrient deficiency stresses were applied to 10 day-old plants. NaH_2_PO_4_ (0.33 mM) and Fe-NaEDTA (0.04 mM) present in the complete media were omitted in the P- and/or Fe-deficient media.

### Iron concentration measurement

Arabidopsis seeds were germinated and grown in the control (+Fe+P) media for 7 days, and then transferred to +Fe+P, iron deficient (-Fe+P), phosphate deficient (+Fe-P), or iron and phosphate deficient (-Fe-P) conditions and grown for 7 additional days. Plants were harvested and shoot samples were dried at 70 °C for 3 days. Total iron was extracted by acid digestion in 1N nitric acid using MARSX (CEM) microwave digester. A 1:10 dilution of the digested material was used to quantify total iron with inductively coupled plasma-atomic emission spectrometry (ICP-OES).

### Analysis of photosystem II activity

Photosystem II (PSII) activity was defined as the maximum quantum yield of the primary quinone acceptor PSII, which was estimated by the ratio of variable fluorescence (Fv) and maximal fluorescence (Fm) of the chlorophyll, Fv/Fm^25^. Arabidopsis wild type (Col-0) seeds were germinated and grown in control (+Fe+P) for 7 days then transferred to three different media: +Fe+P, iron deficient (-Fe+P), and iron and phosphate deficient (-Fe-P) conditions for 0 h (time of the transfer), 15 h, 28 h, 39 h, 52 h, 76 h, 100 h, 124 h, 148 h, and 172 h. Plates containing the seedlings were dark adapted for 30 minutes followed by a very short (160 μs) exposure to a blue measuring beam to determine the minimal fluorescence (F0). The intensity of the detecting and the continuous illumination used was of 156 μE m^-2^s^-1^. A saturating light flash (2600 μE m^-2^s^-1^, 250 ms) was applied to measure the maximum fluorescence (Fm). Kinetics were normalized to the maximum fluorescence (Fm). The maximum quantum yield of Photosystem II (Fv/Fm = (Fm - F0)/Fm) was measured for each growth condition^25^.

### Chlorophyll content measurement

Seeds of Arabidopsis genotypes were germinated and grown in control (+Fe+P) media for 7 days then transferred to three different media: +Fe+P, iron deficient (-Fe+P), and iron and phosphate deficient (-Fe-P) conditions. Fresh leaves (~30mg) were incubated in 2.5mL of 80% acetone overnight in the dark at 4°C. Total chlorophyll content was measured using a UV-VIS spectrophotometer (Beckman Coulter, DU 530). The absorbance of the supernatant was measured at 645 nm and 633 nm. The concentration of total chlorophyll, chlorophyll a, and chlorophyll b were calculated as described previously^26^.

### Ascorbic acid content determination

Seeds of Arabidopsis genotypes were germinated and grown in control (+Fe+P) media and then transferred to +Fe+P, -Fe+P, or -Fe-P media for 76h. Ascorbic acid (AsA) content was measured by a colorimetric assay as described previously^27^. Briefly, shoots were collected and homogenized in ice-cold 6% trichloroacetic acid (TCA) (Sigma Aldrich). In the supernatant, Fe3+ (ferric ion) is reduced by AsA to the Fe2+ (ferrous ion) that, when coupled with 2,2-dipyridyl, forms a complex with a characteristic absorbance at 525 nm. A standard curve was generated using known concentrations of AsA made in 6% TCA to determine the AsA concentration. Blanks were prepared using only 6% TCA. AsA concentration was expressed as μmol g^-1^ fresh weight.

### Hydrogen peroxide quantification

Seeds of Arabidopsis genotypes were germinated and grown in control (+Fe+P) media and then transferred to +Fe+P, -Fe+P, or -Fe-P media for 76h. Hydrogen peroxide (H_2_O_2_) (Sigma-Aldrich) was quantified as described previously^28,29^. Fresh shoot tissues (0.2g) were homogenized with 0.1% (w/v) TCA and were centrifuged at 12,000 g for 15 min at 4 °C. 0.5 ml of supernatant was added to 0.5 ml of 10 mM potassium phosphate buffer (pH 7.0) and 1 ml of 1 M potassium iodide. The absorbance of the reaction mixture was measured at 390 nm. The amount of H_2_O_2_ was calculated using a standard curve prepared from known concentrations of H_2_O_2_ ranging from 0.1 to 1 mM.

### RNA-seq experiments

Arabidopsis wild type (Col-0) plants were grown in control (+Fe+P) media for 7 days and transferred to three different media: control (+Fe+P), iron deficiency (-Fe+P), and iron and phosphate deficiency (-Fe-P) conditions. Shoots were collected at 39h, 52h and 76h after the transfer. For RNA-seq experiments, three biological replicates were prepared for each time point (39h, 52h and 76h) and each condition (+Fe+P, -Fe+P and -Fe-P) for a total of 27 samples. Total RNA was extracted from these samples using RNeasy Plant Mini Kit (QIAGEN) using the RLT buffer supplemented with 2-mercaptoethanol, and RNA quality was verified using an Agilent 2100 BioAnalyzer. The mRNAs were subsequently isolated using magnetic KAPA Biosystems oligo-dT beads from KAPA Biosystems (Roche) and then used for library construction using the KAPA Biosystems RNA HyperPrep Kit (Roche). To index the libraries, we used adapters from the KAPA Biosystems Single-Indexed Adapter Set A+B (Roche). Before pooling the libraries, we monitored their quality and concentrations using an Agilent 2100 BioAnalyzer, Qubit dsDNA HS Assay Kit (Thermo Fisher Scientific) and the KAPA Library Quantification Kit (Roche). Pooled libraries were then sequenced using the NextSeq 500 System at the Stanford Functional Genomics Facility (Stanford, CA). Raw reads were demultiplexed and aligned to the TAIR10 genome annotation using HISAT2^30^ on the Galaxy web platform^31^. Finally, mapped read counts were used to perform normalization and a differential expression analysis on R using the DESeq2^32^ and TxDB.Athaliana.BioMart.plantsmart^28^ (Bioconductor) packages. In DESeq2, p-values from the Wald test were corrected for multiple hypothesis testing using the Benjamini and Hochberg method. A transcript was considered differentially expressed if the adjusted p-value < 0.05. Volcano plots were generated using the EnhancedVolcano package (version 1.6.0) (Bioconductor) with a default cut-off of log2(Fold Change) > |2| and adjusted p value < 10e^-6^. DEGs having a p-value of 0 were converted to 10^-1^x lowest non-zero p-value.

### Real-time quantitative reverse-transcription PCR

Seeds of Arabidopsis wild type (Col-0), *bzip58*, and *pht4;4* mutant plants were germinated and grown for 7 days in control (+Fe+P) media, and then transferred to +Fe+P, Fe+P, or -Fe-P. Shoot tissues were collected at 76h after the transfer, and then used for total RNA extraction as described in^33^. Each experiment was conducted with 16 plants and 4-6 plants were pooled for RNA extraction, resulting in 3-4 biological replicates. Two μg of the total RNA was used for reverse transcription (Promega) to synthesize cDNA using oligo(dT) primer (Promega). Real-time quantitative reverse-transcription PCR (qRT-PCR) was performed as described in^33^ using 384-well plates with a LightCycler 480 Real-Time PCR System (Roche diagnostics). The *Ubiquitin 10* mRNA (*UBQ10:* At4g05320) was used as control to calculate the relative mRNA level of each gene. The primers used in this study are listed in Table S2.

### Genome wide association studies (GWAS)

Gene expression data of the 32 genes that were specifically downregulated by -Fe+P but not by -Fe-P relative to +Fe+P were downloaded from leaf expression data of 727 Arabidopsis accessions ^15^. Normalized RNA-seq read counts of these genes were used to perform Principal Component Analysis, and contributions of the accessions to PC1 that explained 89.5% of the expression variance of the 32 genes were used to run genome-wide association (GWA) analysis. GWA mapping was performed using 1001 genomes SNP data^34^ as implemented in the web application GWAPP^35^. Bonferroni correction (*α* = 0.05) and false-discovery rate (FDR) at 5%^36^ were implemented to account for multiple hypothesis tests.

### Statistical analysis

Box plots were generated using a web based application “BoxPlotR”^37^. Statistical analyses of the data were performed using analysis of variance (ANOVA). One-way ANOVA with a Duncan post-hoc test, and two-way analysis of variance (ANOVA) and Tukey’s honest significant difference (HSD) test were used to compare mean values. For all the statistical analyses, the difference was considered statistically significant when the test yielded a p-value < 0.05.

## Acknowledgements

The authors thank the members of Rhee lab; Benoit Lacombe and HONUDE team (INRAe) for their comments on the manuscript and helpful discussions. We thank the ICP-MS/TIMS Facility within Stanford University for assistance with the ICP-MS measurements, and the Stanford Functional Genomics Facility for assistance with RNA sequencing (Stanford, CA). The GWAS analysis was made possible by data generated by Kawakatsu et al. ^15^.

## Contributions

S.Y.R. and H.R. conceived the project. Experiments were designed by S.Y.R., H.R., H.N., and Z.S. and mainly carried out by H.N. S.C. performed and analyzed experiments related to photosystem II activity. bZIP58-GFP localization, ascorbic acid quantification and hydrogen peroxide assays were conducted by H.C. and N.B. RNA- seq data were generated and analyzed by Y.D. Gene Ontology analysis was performed K.Z. Z.S. performed the genome-wide association mapping. H.R. performed the qRT-PCR analyses, generated plasmid constructs, the homozygote mutants, and the complemented mutant lines. S.Y.R., H.R. and Z.S. wrote the paper with input from all authors.

## Data availability

Data supporting the findings of this work are available within the paper and its Supplementary Information and Source Data files. The datasets and plant materials generated and analyzed during the current study are available from the corresponding author H.R. upon request. Transcriptome data were deposited in NCBI’s Gene Expression Omnibus (GEO) under a project number GSE163190.

## References

1 Myouga, F. et al. The Chloroplast Function Database: a large - scale collection of Arabidopsis Ds/Spm - or T - DNA - tagged homozygous lines for nuclear-encoded chloroplast proteins, and their systematic phenotype analysis. Plant J. 61, 529–542 (2010).

2 Chen, Y. & Barak, P. Iron nutrition of plants in calcareous soils. Adv. Agron. 35, 217–240 (1982).

3 Carstensen, A. et al. The impacts of phosphorus deficiency on the photosynthetic electron transport chain. Plant Physiol. 177, 271–284 (2018).

4 Terry, N. & Abadía, J. Function of iron in chloroplasts. J. Plant Nutr. 9, 609–646 (1986).

5 Terry, N. & Low, G. Leaf chlorophyll content and its relation to the intracellular localization of iron. J. Plant Nutr. 5, 301–310 (1982).

6 Lill, R. Function and biogenesis of iron–sulphur proteins. Nature 460, 831 (2009).

7 Briat, J.-F., Dubos, C. & Gaymard, F. Iron nutrition, biomass production, and plant product quality. Trends Plant Sci. 20, 33–40 (2015).

8 Balk, J. & Pilon, M. Ancient and essential: the assembly of iron–sulfur clusters in plants. Trends Plant Sci. 16, 218–226 (2011).

9 Marschner, H. Mineral nutrition of higher plants. 2nd. (Academic Press, Elsevier, 1995).

10 DeKock, P. C., Hall, A. & Inkson, R. H. E. Active iron in plant leaves. Ann. Bot. 43, 737–740 (1979).

11 Saenchai, C. et al. The involvement of OsPHO1; 1 in the regulation of iron transport through integration of phosphate and zinc deficiency signaling. Front. Plant Sci. 7, 396 (2016).

12 Bournier, M. et al. *Arabidopsis* ferritin 1 *(AtFer1)* gene regulation by the phosphate starvation response 1 (AtPHR1) transcription factor reveals a direct molecular link between iron and phosphate homeostasis. J. Biol. Chem. 288, 22670–22680 (2013).

13 Bertamini, M., Nedunchezhian, N. & Borghi, B. Effect of iron deficiency induced changes on photosynthetic pigments, ribulose-1, 5-bisphosphate carboxylase, and photosystem activities in field grown grapevine *(Vitis vinifera* L. cv. Pinot noir) leaves. Photosynthetica 39, 59–65 (2001).

14 Liu, W. et al. The ethylene response factor AtERF4 negatively regulates the iron deficiency response in *Arabidopsis thaliana*. PLoS ONE 12, e0186580 (2017).

15 Kawakatsu, T. et al. Epigenomic diversity in a global collection of Arabidopsis thaliana accessions. Cell 166, 492–505 (2016).

16 Miyaji, T. et al. AtPHT4; 4 is a chloroplast-localized ascorbate transporter in Arabidopsis. Nat. Commun. 6, 5928 (2015).

17 Torabinejad, J., Donahue, J. L., Gunesekera, B. N., Allen-Daniels, M. J. & Gillaspy, G. E. VTC4 is a bifunctional enzyme that affects myoinositol and ascorbate biosynthesis in plants. Plant Physiol. 150, 951–961 (2009).

18 Noctor, G. & Foyer, C. H. Ascorbate and glutathione: keeping active oxygen under control. Annu. Rev. Plant Biol. 49, 249–279 (1998).

19 Finkel, T. Signal transduction by reactive oxygen species. J. Cell Biol. 194, 7–15 (2011).

20 Shapiguzov, A., Vainonen, J., Wrzaczek, M. & Kangasjärvi, J. ROS-talk–how the apoplast, the chloroplast, and the nucleus get the message through. Front. Plant Sci. 3, 292 (2012).

21 Shaikhali, J. & Wingsle, G. Redox-regulated transcription in plants: emerging concepts. AIMS Mol. Sci. 4, 301 (2017).

22 Warde-Farley, D. et al. The GeneMANIA prediction server: biological network integration for gene prioritization and predicting gene function. Nucleic Acids Res. 38, W214–W220 (2010).

23 Haydon, M. J. et al. Vacuolar nicotianamine has critical and distinct roles under iron deficiency and for zinc sequestration in Arabidopsis. Plant Cell 24, 724–737 (2012).

24 Yoshida, S., Forno, D. A. & Cock, J. H. Laboratory manual for physiological studies of rice. (International Rice Research Institute, 1976).

25 Baker, N. R. Chlorophyll fluorescence: a probe of photosynthesis in vivo. Annu. Rev. Plant Biol. 59, 89–113 (2008).

26 Zhang, J., Han, C. & Liu, Z. Absorption spectrum estimating rice chlorophyll concentration: preliminary investigations. J. Plant Breed. Crop Sci. 1, 223–229 (2009).

27 Gillespie, K. M. & Ainsworth, E. A. Measurement of reduced, oxidized and total ascorbate content in plants. Nat. Protoc. 2, 871–874 (2007).

28 Alexieva, V., Sergiev, I., Mapelli, S. & Karanov, E. The effect of drought and ultraviolet radiation on growth and stress markers in pea and wheat. Plant Cell Environ. 24, 1337–1344 (2001).

29 Gechev, T., Mehterov, N., Denev, I. & Hille, J. A simple and powerful approach for isolation of Arabidopsis mutants with increased tolerance to H2O2-induced cell death. Methods Enzymol. 527, 203–220 (2013).

30 Kim, D., Langmead, B. & Salzberg, S. L. HISAT: a fast spliced aligner with low memory requirements. Nat. Methods 12, 357–360 (2015).

31 Afgan, E. et al. The Galaxy platform for accessible, reproducible and collaborative biomedical analyses: 2018 update. Nucleic Acids Res. 46, W537–W544 (2018).

32 Love, M. I., Huber, W. & Anders, S. Moderated estimation of fold change and dispersion for RNA-seq data with DESeq2. Genome Biol. 15, 550 (2014).

33 Rouached, H. et al. Differential regulation of the expression of two high-affinity sulfate transporters, SULTR1. 1 and SULTR1.2, in Arabidopsis. Plant Physiol. 147, 897–911 (2008).

34 Consortium, O. T. O. G. 1,135 genomes reveal the global pattern of polymorphism in *Arabidopsis thaliana*. Cell 166, 481–491 (2016).

35 Seren, Ü. et al. GWAPP: a web application for genome-wide association mapping in Arabidopsis. Plant Cell 24, 4793–4805 (2012).

36 Benjamini, Y. & Hochberg, Y. Controlling the false discovery rate: a practical and powerful approach to multiple testing. J. R. Stat. Soc. 57, 289–300 (1995).

37 Spitzer, M., Wildenhain, J., Rappsilber, J. & Tyers, M. BoxPlotR: a web tool for generation of box plots. Nat. Methods 11, 121 (2014).

